# MEACA: efficient gene-set interpretation of expression data using mixed models

**DOI:** 10.1101/106781

**Authors:** Bin Zhuo, Duo Jiang

**Keywords:** MEACA, gene-set analysis, pathway, enrichment test, differential expression, gene expression

## Abstract

Competitive gene-set analysis, or enrichment analysis, is widely used for functional interpretation of gene expression data. It tests a known category (e.g. pathway) of genes for enriched differential expression signals. Current methods do not properly capture inter-gene correlations and heterogeneity, resulting in mis-calibration and power loss. We propose MEACA, a new gene-set method based on mixed-effects models. MEACA flexibly incorporates unknown heterogeneity and correlations across genes, and does not need time-consuming permutations. Compared to existing methods, MEACA substantially improves type 1 error control and power in widely ranging scenarios. Real data applications demonstrate MEACA’s ability to recover biologically meaningful relationships.

## Background

Advancements in high-throughput technologies such as microarray and RNA-Seq have made genome-wide expression profiling a popular research tool to study how gene expression patterns associate with experimental, environmental or clinical conditions. A key task of gene expression data analysis involves the detection of differentially expressed genes, which refer to genes whose expression levels are associated with a factor of interest. To this end, the conventional strategy has been to analyze individual genes separately. However, the results from such single-gene analysis are often challenging to interpret, due to the large numbers of genes that are profiled out of which a long list may be significantly differential. To overcome this, a widely used approach has been to study biologically interpretable sets of genes rather than individual genes. Typically, a gene set consists of genes sharing a common biological property (e.g. genes in a known pathway or annotated with a common biological function), and is available through publicly accessible databases such as the Kyoto Encyclopedia of Genes and Genomes (KEGG) [1] and Gene Ontology (GO) [2]. *Gene-set analysis* of gene expression data aims to evaluate the association between the expression levels of genes in a pre-defined set, referred to as *the test set*, and experimental or environmental factors of interest. It examines whether the test set contains or is enriched with differential expression (DE) signals, where the DE signal of a gene can be quantified by comparing the gene’s expression levels across samples grouped according to the factor of interest (e.g. between diseased subjects and healthy controls). Gene-set tests help researchers understand the underlying biological processes in terms of ensembles of genes.

Depending on the null hypothesis that is tested, there are two types of gene-set tests [3]: *self-contained* tests and *competitive* tests (also called *enrichment tests* in some literature). A self-contained test examines the DE signals of genes in the test set without reference to other genes in the genome, with the null being that no genes in the test set are differentially expressed [4, 5, 6, 7, 8, 9]. A competitive test compares DE signals of genes in the test set to those of the genes not in the test set, trying to detect whether the former are more abundant and/or profound than the latter [10, 11, 12]. Many competitive testing methods perform a three-stage analysis [13]. At the first stage, a *gene-level statistic* is calculated for each gene in the whole genome to measure the association between its expression levels and the design variable(s) of interest; such gene-level statistics include, among others, signal-to-noise ratio [14], ordinary *t*-statistic [10] or moderated *t*-statistic [15], log fold change [16] and *z*-score [17]. At the second stage, a set-level statistic is calculated by comparing the gene-level statistics to the genes’ memberships with respect to the test set (i.e., whether a gene belongs to the test set). Examples of the set-level statistics are the enrichment score [14], the maxmean statistic [18], and a statistic derived from convoluted distribution of gene-level statistics [12], to name a few. At the third stage, a *p*-value is obtained for the test set by comparing the set-level statistic to its reference distribution. Compared with self-contained tests, competitive gene-set tests are much more widely used in the genomic literature [19, 11] and will be the focus of our work.

Most competitive gene-set tests assume independence between gene-level statistics [20]. Given that the gene-level statistics are calculated based on a common set of samples, this assumption implicitly requires that the expression levels of different genes are independent. Examples of independence-assuming gene-set tests include, among many others, PAGE [16], the contingency-table-based tests (see [21] for a review) and sigPathway [15, 10]. However, inter-gene correlations can be widespread, for example, among co-regulated genes [19]. It has been recognized that even mild inter-gene correlations may result in severely inflated false positive rate for independence-assuming gene-set tests [18, 19, 3, 11, 12].

A handful of methods have been proposed to account for inter-gene correlations in competitive gene-set tests. One attempt is to evaluate the null distribution of the set-level statistic by permuting the clinical/treatment labels of the samples. Examples include the widely used Gene Set Enrichment Analysis (GSEA) proposed by Subramanian et al. [14] and various other methods [22, 23]. Permuting sample labels does not require an explicit understanding of the underlying correlation structure among genes and thus protects the test against such correlations. Since permuting sample labels is computationally inefficient, Zhou et al. [24] proposed an analytic approximation to permutations for set-level score statistics, which preserves the essence of permutation gene-set analysis with greatly reduced computational burden. However, permuting sample labels in these methods inevitably alters the null and alternative hypotheses of a competitive gene-set test by excluding from the null the possibility that DE signals are present but not enriched in the test set, and consequently confuses the competitive test with the self-contained test, making the results hard to interpret [3, 13, 11].

Another attempt to account for correlated genes is to use set-level statistics that directly incorporate inter-gene correlations estimated from the data. For example, CAMERA [11] calculates a variance inflation factor (VIF) from the sample correlations (after the treatment effects removed) of the observed expression data, which is then incorporated into the set-level statistic to account for the correlations between the gene-level statistics. QuSAGE [12], a recent extension to CAMERA that quantifies gene-set activity with a probability density function, uses a similar VIF to handle inter-gene correlations. MAST [25] adapts CAMERA to the analysis of single-cell RNA-Seq data. However, as we will demonstrate both theoretically and empirically, the VIF approach implicitly assumes that all genes are homogeneous in terms of whether DE is present and the magnitude of the DE effect. This is problematic because DE heterogeneity commonly arises in gene expression studies: in most real data sets, one expects some of the genes to be differentially expressed while others not, and those that are differentially expressed to have varying DE effects. As a result of its failure to account for this heterogeneity, the VIF approach tries to quantify the correlations among gene-level test statistics (e.g., t-statistics) using the within-treatment-group correlations between the expression levels of different genes. However, the former are often smaller than the latter because, when a fraction of the genes are differentially expressed, their DE effects add to the variability of the gene-level statistics across genes and hence act to dilute the correlation between these statistics. We will show that the VIF approach can lead to severely compromised type 1 error and power in gene-set testing.

To address these challenges, we propose a new framework for competitive gene-set analysis, which we will call MEACA (**M**ixed-effects **E**nrichment **A**nalysis with **C**orrelation **A**djustment). Our idea is motivated by the discrepancy, due to DE heterogeneity, between the within-treatment-group correlation structure of the genes expression levels and the correlation structure among gene-level statistics. Using a mixed-model approach, we model the covariance structure of gene-level statistics by two components, one attributable to the correlations between the expression levels of different genes after treatment effects are removed, and the other attributable to the variability across genes in terms of the presence of DE and the effect size. Our method is able to adjust for completely unknown, unstructured correlations among the genes. We use a quasi-likelihood framework, which does not require the gene expression data or the distribution of the DE effects across genes to be Gaussian. MEACA uses a score-type test and allows for analytical assessment of *p*-values, which renders it computationally efficient for analysis of large numbers of genes and gene sets. We will show that, compared to existing methods including GSEA [14] and CAMERA [11], MEACA consistently outperforms existing methods in terms of type 1 error control in a wide variety of correlation settings and enjoys substantial power gain.

## Results

### Method Overview

In enrichment analysis, we are interested in a pre-defined set of genes, for example, from a known pathway or given by a functional annotation term from a database such as KEGG [1] or GO [2]. Our goal is to test whether this known gene set is enriched with DE signals compared to the rest of the genes. We will refer to the genes in the pre-defined gene set as “the test genes” which make up the “the test set,” and genes not in this set “the background genes” which make up “the background set.”

Our gene-set testing method, MEACA, is based on a mixed-effects model that flexibly incorporates the unknown distribution of DE effects and effectively adjusts for completely unknown, unstructured correlations among genes. MEACA does not rely on time-consuming permutations, and uses a quasi-likelihood framework which does not assume the data follows a normal distribution. Our methodology is connected to that of CAMERA, in that both estimate an inter-gene correlation matrix from the data to adjust the distribution of the gene-level statistics. However, MEACA uses a modeling approach that does not rely on either of the following two assumptions, which are implicitly required by CAMERA but are likely violated in reality:

(A1) Either none of the genes are differentially expressed or all genes are differentially expressed with the exact same DE effect, both in the test set and in the background set;
(A2) Inter-gene correlations are present only among genes in the test set, not among background genes or between background and test genes.

More details on these assumptions and the MEACA methodology will be presented in Methods.

### Simulation Studies

We conduct type 1 error and power simulations to evaluate the performance of MEACA and compare it with other methods. In this section, we outline the essential components of these simulations, with additional details provided in Methods. First, to assess the impact of DE heterogeneity on the performance of various methods, we conduct two groups of simulations: In group I simulations, no background genes are differentially expressed, so under the null hypothesis no DE genes are present either in the test set or the background set; In group II simulations, a proportion of background genes are allowed to be differentially expressed, and under the null the same proportion are differentially expressed among test genes. Hence, Assumption (A1) holds for group I but not for group II. Let *p_b_* and *p_t_* be the DE probabilities for a background gene and a test gene, respectively. Table 1 summarizes the configurations of *p_b_* and *p_t_* we consider, in both type 1 error (configuration *S*_null_) and power simulations (configurations *S*_1_ − *S*_4_).

**Table 1.** DE probability configurations in type 1 error and power simulations.

| Group | Background | DE prob. in test set ($p_t$) | | | | |
| --- | --- | --- | --- | --- | --- | --- |
| | DE prob. ($p_b$) | $S_{\text{null}}$ | $S_1$ | $S_2$ | $S_3$ | $S_4$ |
| I | 0% | 0% | 5% | 10% | 15% | 20% |
| II | 10% | 10% | 15% | 20% | 25% | 30% |

To study how inter-gene correlation affects the methods to be compared, we examine five different correlation structures, (a)-(e). We assume a common pairwise correlation coefficient for genes from the same category (either the test set or the background set): let *ρ*_1_ be the correlation between test genes, *ρ*_2_ be the correlation between background genes, and *ρ*_3_ be the correlation between a test gene and a background gene. We examine five different correlation structures, listed as follows:

a. *ρ*_1_ = *ρ*_2_ = *ρ*_3_ = 0; that is, the genes are independent of each other.
b. *ρ*_1_ = *ρ*_2_ = *ρ*_3_ =0.1; that is, all genes are correlated, with an exchangeable correlation structure.
c. *ρ*_1_ = 0.1, *ρ*_2_ = *ρ*_3_ = 0; that is, only the genes in the test set are correlated.
d. *ρ*_1_ = 0.1, *ρ*_2_ = 0.05, *ρ*_3_ = 0; that is, genes are correlated within the test set and within the background set, but any two genes, one from the test set and the other from the background set, are independent.
e. *ρ*_1_ = 0.1, *ρ*_2_ = 0.05, *ρ*_3_ = −0.05; that is, all genes are correlated, but the correlation between two genes depends on their membership status to the test set.

The five structures will help us evaluate the robustness of MEACA and how violations of the independence assumption or Assumption (A2) affect the competing methods.

In the simulations, we compare MEACA to five existing gene-set testing methods: sigPathway [10], MRGSE [26], CAMERA [11], QuSAGE [12], and GSEA [14]. MRGSE is a rank-based method assuming inter-gene independence, and is recommended by Tarca et al. [27] as the best performing one among a wide class of independence-assuming methods. sigPathway is a parametric version of MRGSE, and in our simulations we use the moderated *t*-statistic [15] as its gene-level statistic. The other three methods in comparison, CAMERA, QuSAGE [12], and GSEA [14], all incorporate features intended for inter-gene correlation correction. CAMERA uses the moderated *t*-statistic as the gene-level statistic and estimates a VIF to account for inter-gene correlations. QuSAGE is an extension of CAMERA that quantifies gene-set activity with a probability density function. GSEA first calculates an enrichment score for a test set from the ranks of all genes based on DE evidence, and then determines the significance of the enrichment score by randomly permuting the case-control labels of the samples.

#### Type 1 error simulations

We evaluate the calibration of MEACA and the competing methods using data simulated under a variety of settings. For MEACA and five other approaches (sigPathway, MRGSE, GSEA, CAMERA and QuSAGE), Figure 1 shows the quantile-quantile (QQ) plots of *p*-values in simulation groups I (left column) and II (right column) and under each of the five correlation structures (each row, from top to bottom, corresponds accordingly to correlation structures (a)-(e)). The plots are based on 10,000 simulation replicates. In each QQ plot, the vertical axis corresponds to the empirical *p*-values and the horizontal axis corresponds to quantiles from the uniform distribution between 0 and 1, which is the theoretical distribution of the *p*-values if a method is correctly calibrated. For any given setting, a curve that closely follows the diagonal line indicates a method that is well calibrated. A curve that falls consistently below the diagonal line indicates a method that has inflated type 1 error, whereas a curve consistently above the diagonal line indicates overly conservative type 1 error control.

**Figure 1.**
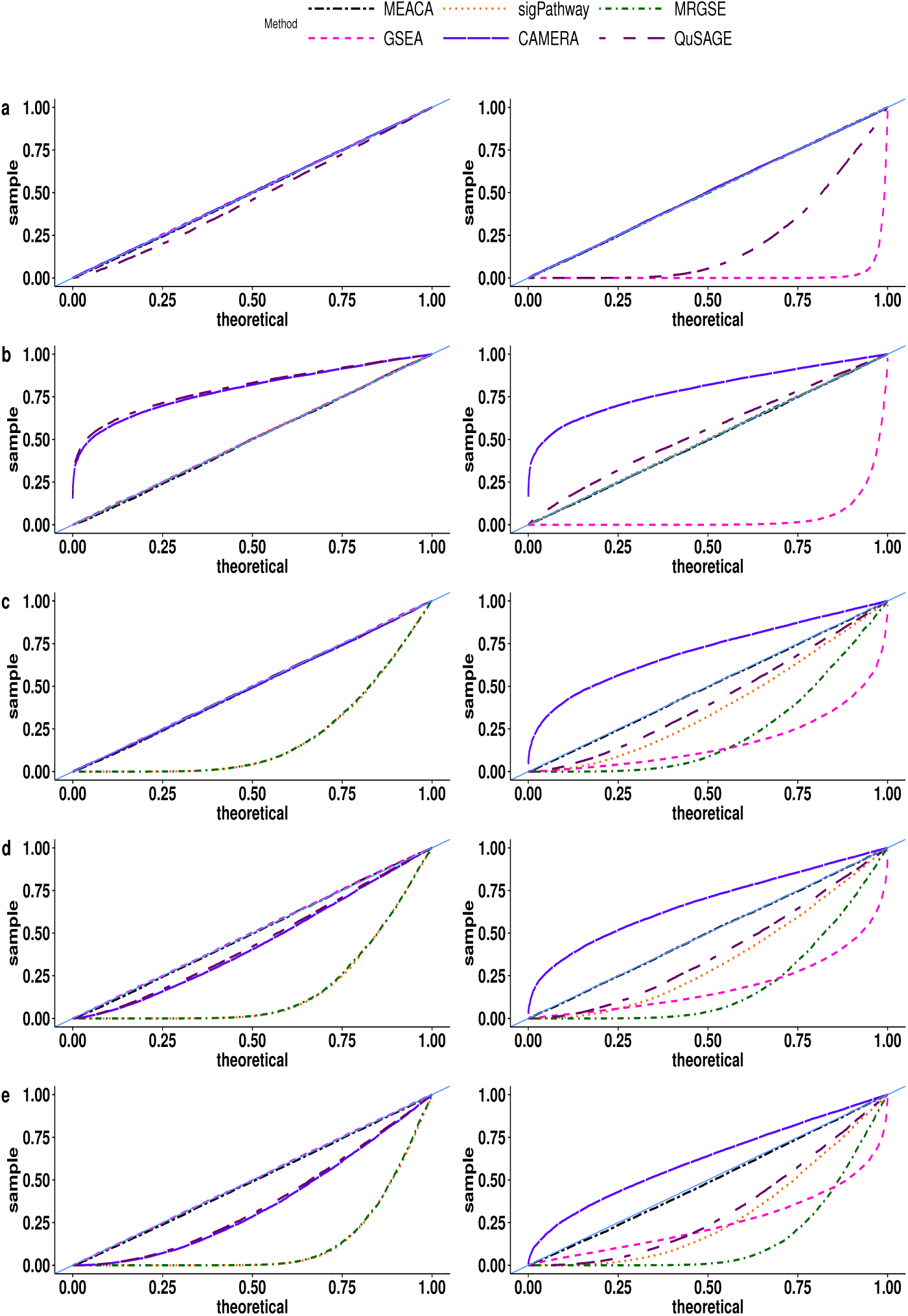
Quantile-quantile plots for *p*-values by different methods in type 1 error simulations. The plots from top to bottom correspond to the correlation structures (a)-(e), respectively. The left column is for group I simulation, and the right column for group II simulation (see Table 1 for details). Results are based on 10, 000 simulation replicates. MEACA gives uniformly distributed *p*-values under all simulation settings, whereas all of the other methods can be severely mis-calibrated under some settings.

Across the settings, MEACA shows consistent accuracy for type 1 error control. All the other methods, however, can be severely mis-calibrated under various scenarios. In particular, the independence-assuming methods, sigPathway and MRGSE, are well calibrated only when the genes are uncorrelated (structure (a)) or when the genes are equally correlated (structure (b)). When the genes go beyond these simple structures, sigPathway and MRGSE become very liberal ((c)-(e))), with type 1 error rates at level 0.05 as high as 0.68 (structure (e), group I). These results show that even small inter-gene correlations (e.g. 0.05) can result in inflated type 1 error if the test does not account for such correlations.

For GSEA, accuracy of type 1 error control relies on the absence of background DE signals: in group I where no gene is differentially expressed, GSEA performs extremely well; group II, however, reveals the failure of GSEA in controlling type 1 error when DE signals are present in both the test and the background sets, regardless of whether inter-gene correlations exist or not. This phenomenon is not surprising given that GSEA permutes the case-control labels of samples, which inevitably disturbs the DE patterns in the genes and is effectively testing a very restrictive null hypothesis, one in which not only the set of test genes cannot be enriched with DE signals compared to the set of background genes, but in fact neither set is allowed to contain any differentially expressed genes at all. This null hypothesis implies the null entailed by the goal of competitive gene-set testing, which is why GSEA is correct in group I. But the former is much more restrictive than the latter, which explains GSEA’s anti-conservativeness in group II. It is notable that, in practice, one rarely sees a situation where no differentially expressed genes are present in the background set, so group II is more relevant than group I, making GSEA a risky choice for the purpose of competitive gene-set testing.

For CAMERA, control of type 1 error varies from being too conservative to being too liberal across the settings in Figure 1. For any given setting, the performance of CAMERA would depend on (1) whether DE effects are heterogeneous across genes and (2) the inter-gene correlation structure. These two factors correspond, respectively, to Assumptions (A1) and (A2) discussed in Material & Methods. In simulation group I, (A1) holds because DE effects are completely absent and therefore homogeneous across genes. In this case, CAMERA is correctly calibrated under (a) and (c), when Assumption (A2) also holds. When all genes, including the background genes, are correlated (as is the case for structure (b)), CAMERA is overly conservative with type 1 error rate at level 0.05 too stringently controlled at < 10^−4^. Under structures (d) and (e), CAMERA tends to be too liberal, with type 1 error at level 0.05 as high as 0.21 (structure (e), group I). QuSAGE has similar trends of mis-calibration in these group I settings, and is anti-conservative under (a). In contrast to group I, group II has a fraction of the genes that are differentially expressed with varying effects, resulting in heterogeneity among genes in terms of the presence and magnitude of DE effects. So in this case Assumption (A1) is violated. As discussed in Material & Methods, this would drive the type 1 error of CAMERA towards the conservative side when inter-gene correlation is present, because CAMERA ignores the DE heterogeneity and consequently would over-correct for the correlation. Indeed, as shown in the right column of Figure 1, when genes are correlated (structures (b)-(e)), the calibration of CAMERA is very conservative, with type 1 error at level 0.05 falling below 0.005. Such stringent control of type 1 error is expected to come at the cost of low power of detecting gene sets that are truly enriched with DE signals, which we will show in the power simulations. In group II, QuSAGE is also mis-calibrated across the settings.

#### Power simulations

Figure 2 shows how the power of MEACA varies as the enrichment in the test set becomes more profound (from *S*_1_ to *S*_4_) in the alternative hypothesis. For each correlation structure, we report the power trajectory at level 0.05. The top is the power for group I, and the bottom for group II. The power results under correlation structures (a) and (b) are similar, and are among the highest under each of the four alternatives. As the correlation structure becomes more complex, from (c) to (d) then to (e), the power decreases under every alternative setting. The power under correlation structure (e) is the lowest for both groups I and II.

**Figure 2.**
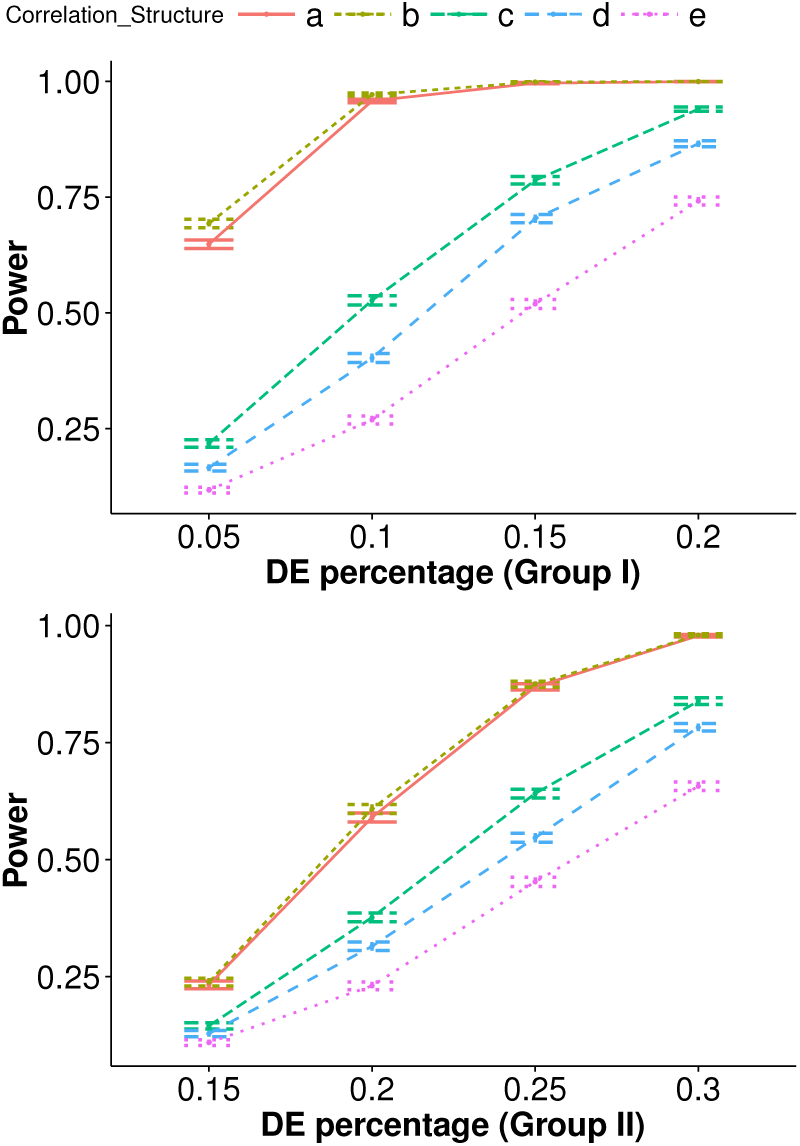
Power of MEACA under correlation structures (a)-(e). The top corresponds to group I simulations, and the bottom to group II simulations (see Table 1). The error bars are the 95% confidence intervals based on 10,000 simulation replicates.

It is also of interest to explore whether MEACA, while being able to adjust for inter-gene correlations, will have compromised power when genes are in fact uncorrelated. For this purpose, we compare the empirical power of MEACA, MRGSE, sig-Pathway and CAMERA under correlation structure (a). We do not consider GSEA or QuSAGE because they do not have consistently accurate control of type 1 error under (a). In Table 2, it is clear that MEACA does not lose any power compared to the independence-assuming methods when the genes are indeed independent. CAMERA also loses little power under structure (a). However, we note that, in the presence of inter-gene correlations, CAMERA is expected to lose power in many realistic scenarios due to its over-stringent calibration (Table 3), and independence-assuming methods tend to generate excessive false positives (Figure 1).

**Table 2.** Power comparison under (a), when genes are *uncorrelated*.

| Group | Method | $S_1$ | $S_2$ | $S_3$ | $S_4$ |
| --- | --- | --- | --- | --- | --- |
| I | MEACA | **0.65** | **0.96** | **1.00** | **1.00** |
|  | CAMERA | 0.63 | 0.95 | **1.00** | **1.00** |
|  | MRGSE | 0.12 | 0.31 | 0.58 | 0.80 |
|  | sigPathway | **0.65** | **0.96** | **1.00** | **1.00** |
| II | MEACA | **0.23** | **0.59** | **0.87** | **0.98** |
|  | CAMERA | **0.23** | 0.58 | 0.86 | **0.98** |
|  | MRGSE | 0.11 | 0.31 | 0.58 | 0.83 |
|  | sigPathway | **0.23** | **0.59** | **0.87** | **0.98** |

MEACA, while being able to account for inter-gene correlations, does not lead to power loss when the genes are in fact uncorrelated. Empirical power at level 0.05 is calculated for each of the four alternative settings $S_1$ - $S_4$ and groups I and II (see Table 1 for details). Results are based on 10,000 simulation replicates. The highest power is in bold type for each setting.

**Table 3.** Power comparison under correlation structures (a)-(e) for group II.

| Method | (a) | (b) | (c) | (d) | (e) |
| --- | --- | --- | --- | --- | --- |
| MEACA | **0.87** | **0.87** | **0.64** | **0.55** | **0.45** |
| CAMERA | 0.86 | 0.028 | 0.061 | 0.076 | 0.11 |
| MRGSE | 0.58 | 0.58 | — | — | — |
| sigPathway | **0.87** | **0.87** | — | — | — |
| GSEA | — | — | — | — | — |
| QuSAGE | — | 0.84 | — | — | — |

Finally, we compare the statistical power of MEACA to the other methods under different correlation structures (Table 3). Note that it is not fair or interesting to compare to a method that does not effectively control false positives. Therefore, to make a more meaningful power comparison, for any given setting, we only consider the methods whose type 1 error control is adequate (i.e. either accurate or conservative, but not anti-conservative) as shown by Figure 1, and we leave out a method if its type 1 error rate is inflated. For example, in group II under correlation structure (c), all of MRGSE, sigPathway, GSEA and QuSAGE are anti-conservative and therefore excluded, whereas we include CAMERA which is conservative and MEACA which is accurate. We focus on the group II scenarios, which we consider more practically relevant than group I because in real data sets one typically expects at least some of the background genes to be differentially expressed. Table 3 shows that MEACA enjoys the highest power under all of the correlation structures. CAMERA is the only other method that is adequately calibrated across the settings. However, CAMERA has by far a lower power when the genes are correlated, with the power at level 0.05 as low as 0.028 (structure (b)). This aligns with the highly conservative type 1 error control of CAMERA when DE signal is present among the background genes (Figure 1). Our results indicate that MEACA consistently maintains the highest power and achieves great power gain over CAMERA, which can be greatly underpowered in some realistic settings.

### Real Data

We conduct competitive gene-set analysis on two real data sets to illustrate the use of MEACA and to compare the enriched gene sets it identifies with those obtained by three other methods, GSEA, CAMERA and MRGSE.

#### Huntington’s Disease Data

We examine an RNA-Seq data set on the Huntington’s Disease (HD) to identify enriched gene sets that are potentially responsible for HD. The mRNA expression profiles in human prefrontal cortex were obtained from 20 Huntington’s Disease samples and 49 neurologically normal controls. Expression values were normalized and filtered as described in [28]. The data set, containing 28, 087 genes is available as series GSE64810 in the GEO database (http://www.ncbi.nlm.nih.gov/geo/). For each gene, we adjust for two covariates—age at death (DeathAge) and RNA Integrity Number (RIN), both treated as categorical variables [28]. Briefly, DeathAge is binned into intervals 0-45, 46-60, 61-75, 76-90 and 90+, and RIN is dichotomized as > or ≤ 7. We regress the normalized expression levels on AgeDeath and RIN and use the resulting residuals as the covariate-adjusted expression levels.

We perform enrichment analysis on the covariate-adjusted data using the MsigDB [14] C2 Canonical Pathways (February 5, 2016, data last accessed). The C2 Canonical Pathways have a collection of 1330 gene sets, with an average size of 50 genes (the size ranges from 3 to 1028, and the median is 29). Since the genes are named by HGNC symbols in C2 and by Ensembl IDs in the HD expression data set, we convert the Ensembl IDs in the expression data into HGNC symbols using *BioMart* (http://uswest.ensembl.org/biomart/martview/). We retain 26, 941 genes that have corresponding HGNC symbols. On each gene set in the entire collection of C2 Canonical Pathways, we perform four testing methods (MEACA, GSEA, CAMERA and MRGSE) to obtain *p*-values evaluating whether the gene set is enriched with DE signals associated with HD.

In Figure 3 we plot the *p*-values of MEACA against those of GSEA, CAMERA and MRGSE on the negative *log*10 scale. The *p*-values of CAMERA are overwhelmingly larger than those of GSEA and MEACA, yet smaller than those of MRGSE. This is consistent with our observation in the type 1 error simulations that CAMERA can produce conservative *p*-values. The *p*-values of MEACA and those of the other three methods are highly correlated (Pearson’s correlations of log 10 *p* between MEACA and GSEA, CAMERA and MRGSE are 0.91, 0.96, and 0.81, respectively). The *p*-values of MRGSE are in general smaller than the corresponding *p*-values of MEACA, likely due to unadjusted inter-gene correlations and leading to more gene sets claimed to be significant by MRGSE.

**Figure 3.**
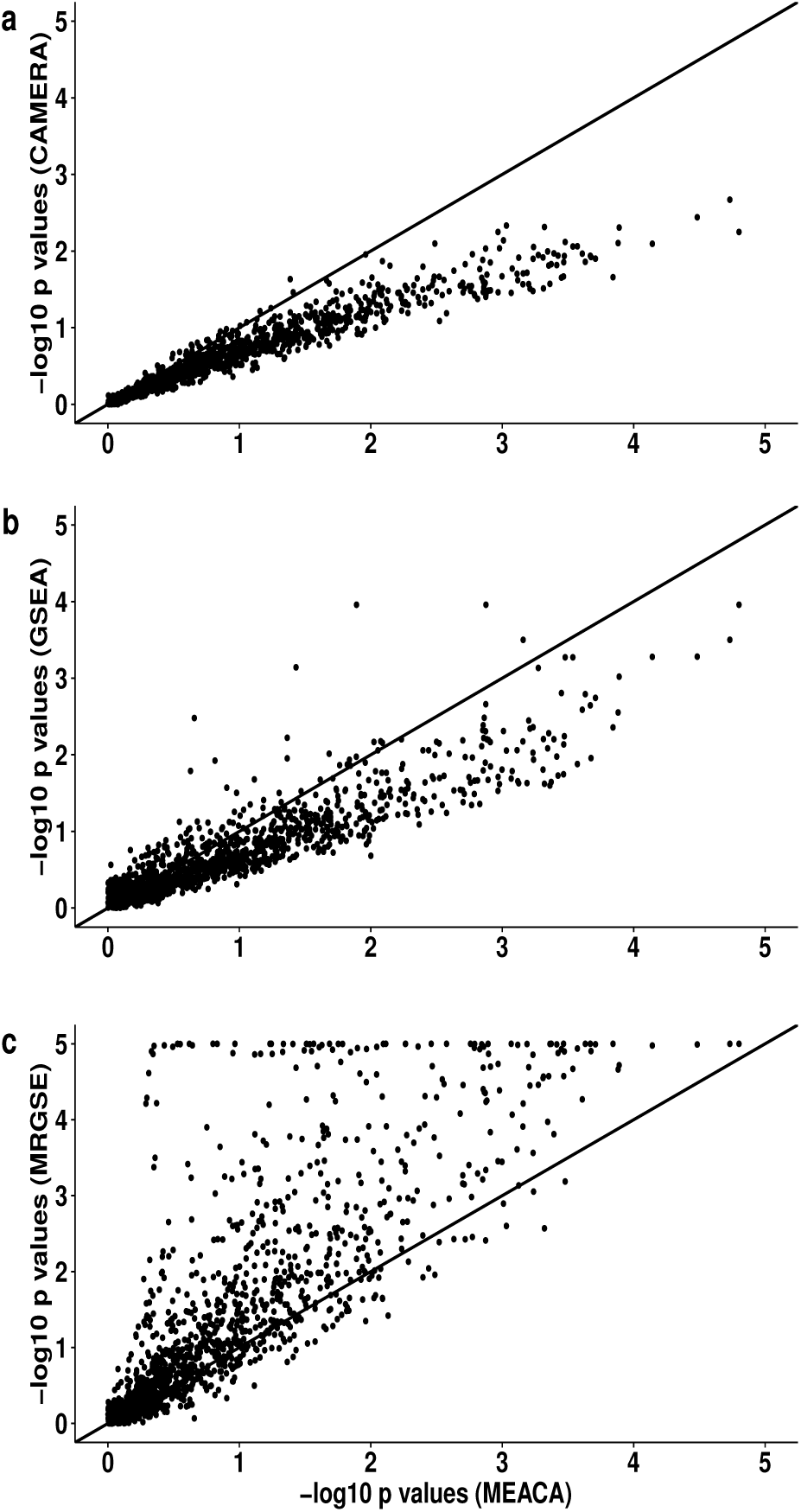
Comparison of *p*-values between MEACA, GSEA, CAMERA and MRGSE in HD data. The *p*-values are reported from enrichment test of each gene set in the C2 Canonical Pathway gene sets.

We then compare the resulting list of significant gene sets identified by MEACA to those by the other three methods. For multiple comparison adjustment, we use the Benjamini-Hochberg [29] procedure (BH) to control the false discovery rate (FDR) at 0.05. Out of a total of 1330 C2 Canonical Pathways, MEACA identifies 89 gene sets to be significantly enriched. In contrast, GSEA identifies 3 enriched gene sets—2 of them are also among those 89 gene sets identified by MEACA (the one that is not significant according to MEACA has a nominal *p*-value of 0.013 and a BH-adjusted *p*-value of 0.100). MRGSE identifies as many as 371 gene sets, which include all the 89 sets identified by MEACA as well as 282 other gene sets, which are likely to contain many false discoveries due to MRGSE’s failure to control for inter-gene correlations. CAMERA fails to detect any significant gene set. In their original paper, Labadorf et al. [28] used the same HD data set to conduct enrichment analysis with topGo [30]. They noted that the enriched gene sets they identified showed a clear immune response and inflammation-related pattern, including the PID NF-kappaB canonical pathway, PID IL4-mediated signaling events (Pathway name: PID_IL4_2PATHWAY) and the Reactome innate immune system pathway. In our analysis, MEACA is able to capture all of these three gene sets, which rank (by nominal *p*-values) 3, 10 an 18, respectively, among the 89 enriched gene sets by MEACA.

In Table 4, we report the top 30 enriched gene sets (ordered by nominal *p*-values) identified using MEACA. Among these, only one gene set (labeled by “*” in the table) is also identified by GSEA at FDR level of 0.05, and none by CAMERA.

**Table 4.** Top 30 enriched gene sets identied by MEACA for HD data.

| Gene Set | Size | $\hat{\rho}_1$ | $\hat{\rho}_2$ | $\hat{\rho}_3$ | $p$-value | BH-adjusted $p$-value |
| --- | --- | --- | --- | --- | --- | --- |
| PID SMAD2 3NUCLEAR PATHWAY | 79 | 0.063 | 0.013 | 0.015 | 5.8E-06 | 5.7E-03 * |
| REACTOME YAP1 AND WWTR1 TAZ STIMULATED GENE EXPRESSION | 23 | 0.121 | 0.013 | 0.014 | 8.5E-06 | 5.7E-03 |
| PID NFKAPPAB CANONICAL PATHWAY | 22 | 0.127 | 0.013 | 0.019 | 2.3E-05 | 1.0E-02 |
| BIOCARTA NTHI PATHWAY | 23 | 0.130 | 0.013 | 0.023 | 6.2E-05 | 2.1E-02 |
| BIOCARTA TID PATHWAY | 18 | 0.101 | 0.013 | 0.012 | 1.2E-04 | 2.2E-02 |
| PID HIV NEF PATHWAY | 35 | 0.065 | 0.013 | 0.013 | 1.2E-04 | 2.2E-02 |
| KEGG PATHWAYS IN CANCER | 311 | 0.028 | 0.013 | 0.010 | 1.3E-04 | 2.2E-02 |
| PID MYC REPRESS PATHWAY | 60 | 0.057 | 0.013 | 0.013 | 1.9E-04 | 2.2E-02 |
| BIOCARTA TOLL PATHWAY | 36 | 0.083 | 0.013 | 0.018 | 2.0E-04 | 2.2E-02 |
| PID IL4 2PATHWAY | 59 | 0.081 | 0.013 | 0.010 | 2.0E-04 | 2.2E-02 |
| KEGG TGF BETA SIGNALING PATHWAY | 82 | 0.055 | 0.013 | 0.011 | 2.2E-04 | 2.2E-02 |
| BIOCARTA DEATH PATHWAY | 33 | 0.067 | 0.013 | 0.013 | 2.4E-04 | 2.2E-02 |
| KEGG NOD LIKE RECEPTOR SIGNALING PATHWAY | 55 | 0.045 | 0.013 | 0.008 | 2.6E-04 | 2.2E-02 |
| BIOCARTA CTCF PATHWAY | 23 | 0.083 | 0.013 | 0.015 | 2.8E-04 | 2.2E-02 |
| ST TUMOR NECROSIS FACTOR PATHWAY | 28 | 0.031 | 0.013 | 0.014 | 3.2E-04 | 2.2E-02 |
| BIOCARTA TNFR2 PATHWAY | 17 | 0.151 | 0.013 | 0.022 | 3.3E-04 | 2.2E-02 |
| KEGG APOPTOSIS | 82 | 0.036 | 0.013 | 0.008 | 3.3E-04 | 2.2E-02 |
| REACTOME INNATE IMMUNE SYSTEM | 209 | 0.039 | 0.013 | 0.009 | 3.3E-04 | 2.2E-02 |
| PID HES HEY PATHWAY | 47 | 0.071 | 0.013 | 0.019 | 3.4E-04 | 2.2E-02 |
| REACTOME DOWNSTREAM TCR SIGNALING | 31 | 0.082 | 0.013 | 0.011 | 3.7E-04 | 2.2E-02 |
| PID TCPTP PATHWAY | 42 | 0.076 | 0.013 | 0.010 | 3.7E-04 | 2.2E-02 |
| BIOCARTA 41BB PATHWAY | 14 | 0.110 | 0.013 | 0.023 | 3.9E-04 | 2.2E-02 |
| PID FRA PATHWAY | 34 | 0.154 | 0.013 | 0.008 | 4.1E-04 | 2.2E-02 |
| PID P53 DOWNSTREAM PATHWAY | 131 | 0.045 | 0.013 | 0.012 | 4.2E-04 | 2.2E-02 |
| PID EPO PATHWAY | 34 | 0.069 | 0.013 | 0.013 | 4.3E-04 | 2.2E-02 |
| BIOCARTA PPARA PATHWAY | 53 | 0.031 | 0.013 | 0.008 | 4.4E-04 | 2.2E-02 |
| BIOCARTA EPONFKB PATHWAY | 11 | 0.068 | 0.013 | 0.010 | 4.7E-04 | 2.2E-02 |
| BIOCARTA HIVNEF PATHWAY | 58 | 0.063 | 0.013 | 0.019 | 4.8E-04 | 2.2E-02 |
| BIOCARTA CD40 PATHWAY | 13 | 0.165 | 0.013 | 0.026 | 4.8E-04 | 2.2E-02 |
| BIOCARTA IL7 PATHWAY | 17 | 0.100 | 0.013 | 0.016 | 5.2E-04 | 2.3E-02 |

The coefficients, $\hat{\rho}_1$ , $\hat{\rho}_2$ and $\hat{\rho}_3$ , respectively, are the average estimated sample correlations of observed data between genes in the test set, between genes in the background set, and between two genes belonging to two different sets. A gene set significantly enriched by GSEA is indicated by “\*”. No gene set is identified as enriched by CAMERA and all the 30 gene sets are among the 371 genes identified by MRGSE. For all methods, FDR is controlled at 0.05.

The majority of the enriched gene sets by MEACA have been previously shown to be closely related to HD pathogenesis. For example, the top enriched gene set, PID SMAD2 3NUCLEAR PATHWAY, is responsible for regulation of nuclear SMAD2/3 signaling. Nuclear SMAD2/3 has been linked to polyglutamine diseases, a group of neurodegenerative disorders that include HD [31]. The second gene set, REACTOME YAP1 AND WWTR1 TAZ STIMULATED GENE EXPRESSION, consists of genes whose expressions are regulated by transcriptional co-activators YAP1 and WWTR1. YAP1 has been extensively linked to HD [32, 33, 34]. The third enriched gene set, PID NFKAPPAB CANONICAL PATHWAY, is a canonical NF-kappaB pathway, and its dysregulation has been shown on the cellular level to cause HD immune dysfunction [35]. It has also been found that reduced transport of NF-kappaB out of dendritic spines and its activity in neuronal nuclei may contribute to the etiology of HD [36]. This also suggests that the BIOCARTA NTHI PATHWAY, related to NF-kappaB activation, is a plausible pathway associated with HD. Moreover, the PID HIV NEF PATHWAY, is a pathway for negative effector of Fas and TNF-alpha, both of which are proteins that have been linked to HD in mice [37]. Furthermore, three of the enriched gene sets, PID MYC REPRESS PATHWAY, BIOCARTA TOLL PATHWAY, and KEGG NOD LIKE RECEPTOR SIGNALING PATHWAY, involve C-MYC, toll-like receptors and NOD-like receptors, respectively, all of which have previously been found to relate to HD or other neurodegenerative disorders [38, 39, 40]. The KEGG TGF BETA SIGNALING PATHWAY has been associated with HD using an independent data set [41]. Another gene set, REACTOME INNATE IMMUNE SYSTEM, has been found to contribute to HD pathogenesis [28, 35]. In addition, Chiang et al. [42] demonstrated that the systematic downregulation of PPAR*γ*, related to the BIOCARTA PPARA PATHWAY, seems to play a critical role in the dysregulation of energy homeostasis observed in HD, and that PPAR*γ* is a potential therapeutic target for this disease. For PID P53 DOWNSTREAM PATHWAY, Ghose et al. [43] have shown the likely involvement of NFkB (RelA), p53 and miRNAs in the regulation of cell death in HD pathogenesis.

#### Male vs Female Lymphoblastoid Cells Data

As a way to validate our method, we analyze the mRNA expression profiles from lymphoblastoid cell lines derived from 17 females and 15 males. Subramanian et al. [14] examined this data set with their GSEA method, testing the MsigDB cytogenetic gene sets (C1) for association with sex. The C1 collection includes 24 gene sets, one for each of the 24 human chromosomes, and 295 gene sets corresponding to cytogenetic bands. Comparing male and female cell lines, one would expect to home in on gene sets on chromosome Y[14]. Because of this prior knowledge, we use this data set as a benchmarking tool to compare different testing methods.

We perform enrichment analysis with four tests, MEACA, GSEA, CAMERA and MRGSE, on all the 309 C1 gene sets containing at least 3 genes. Again, the GSEA *p*-values are obtained using (*b* + 1)/(*K* + 1) with *K* = 9999. In Table 5, we summarize all the gene sets that are identified to be significant by at least one of the four testing procedures, with FDR controlled at 0.05 by the BH procedure. MEACA has recapitulated our knowledge about the data set to a great extent in that it identifies all and only the four gene sets corresponding to chromosome Y or Y bands. In comparison, GSEA, CAMERA and MRGSE not only yield less significant *p*-valuet than MEACA for three of the gene sets on chromosome Y, but have also missec the fourth gene set, chrYp22. Moreover, MRGSE claims as significant three autosomal chromosomes which do not show evidence of enrichment by any of the othei methods.

**Table 5.** Enriched gene sets and their BH-adjusted *p*-values for lymphoblastoid cells data.

| Gene set | Size | MEACA | GSEA | CAMERA | MRGSE |
| --- | --- | --- | --- | --- | --- |
| chrY | 40 | **<1.0E-15** | **1.1E-02** | **1.6E-03** | **9.3E-05** |
| chrYq11 | 16 | **<1.0E-15** | **1.1E-02** | **2.3E-05** | **8.9E-04** |
| chrYp11 | 18 | **8.5E-13** | **1.1E-02** | **3.0E-02** | **2.7E-02** |
| chrYp22 | 8 | **3.9E-02** | 6.8E-01 | 8.2E-01 | 2.2E-01 |
| chr6 | 614 | 8.7E-01 | 1.0E-00 | 1.0E-00 | **1.3E-02** |
| chr1 | 1104 | 8.7E-01 | 1.0E-00 | 1.0E-00 | **4.2E-03** |
| chr12 | 571 | 8.8E-01 | 1.0E-00 | 1.0E-00 | **1.6E-06** |

Reported are gene sets with BH-adjusted $p$ -value < 0.05 for at least one of MEACA, GSEA, CAMERA and MRGSE. An adjusted $p$ -value is made bold if below 0.05.

## Discussion and Conclusions

We have developed MEACA, a new method for competitive gene-set analysis of gene expression data, with the aim of evaluating the association between a set of genes and a factor of interest. MEACA features effective adjustment for completely unknown, unstructured correlations among the genes, and the ability to account for the DE heterogeneity across genes. It uses a score test approach and allows for analytical assessment of *p*-values without the need of time-consuming permutation procedures. Compared to previously proposed approaches, MEACA enjoys robust and accurate control of type 1 error and maintains high power across a wide range of settings. Our method is available in the `MEACA R` package.

Inter-gene correlations are widespread in gene expression data, and failure to account for such correlations has been extensively shown to be problematic for gene-set analysis. Under the competitive gene-set testing framework, a number of methods have been proposed to account for correlation among genes. One approach is to evaluate the significance of set-level statistic by permuting sample labels, as adopted by the widely used procedure GSEA [14]. However, the sample permutation method has been criticized for altering the null hypotheses being tested in the competitive gene-set analysis [3, 13] and consequently tends to result in mis-calibrated testing results. Instead, CAMERA [11] and a recent extension, QuSAGE [12], correct for the correlations among genes by estimating a VIF directly from the data. We are the first to point out a major problem with this approach related to its failure to properly model the DE heterogeneity across genes, which results in incorrect adjustment for the correlation between single-gene test statistics. We have shown in both simulations and real data examples that this can severely compromise the performance of CAMERA and QuSAGE. In particular, we have found that CAMERA can be profoundly mis-calibrated and underpowered under realistic scenarios. We have addressed this challenge by modeling the covariance structure between gene-level statistics using two variance components, one attributable to correlations between gene expressions after potential DE effects are removed, and the other attributable to the heterogeneity of DE effects. Moreover, MEACA is based on a quasi-likelihood framework, which does not assume normality for the expression data or the distribution of the DE effects.

We have compared the performance of MEACA to competing approaches through both simulations and real data examples. Through extensive simulation studies, we have examined the calibration of MEACA and five other methods (sigPathway, MRGSE, CAMERA, GSEA and QuSAGE) in a variety of settings, and have demonstrated that MEACA controls type 1 error accurately under all settings considered, whereas each of the other methods has failed in at least some situations. The power of MEACA is also shown to compare favorably with the other methods. We have further validated our approach using two real data sets, in which MEACA, compared with its competitors, has yielded results that are highly biologically relevant. In particular, we have identified a moderate number of gene sets associated with HD, many of which have previously been linked to the disease yet most, if not all, of which were missed by GSEA and CAMERA. As a simple benchmarking data set, we have also analyzed a lymphoblastoid cell line data set for which we have relatively confident prior understanding. MEACA has been able to generate results that are highly consistent with our prior knowledge.

Although MEACA is motivated by the problem of gene-set analysis of transcriptomic data, it can be widely applicable to other types of data sets (such as proteomic, metabolomic and microbiome data) in which it is of interest to detect whether a subset of the features (such as protein categories, metabolite groups and microbial taxonomic groups) are enriched with differential signals between two groups of samples. Examples include detection of differentially abundant gene families in functional analysis of metagenomic data [44] and enrichment analysis of high-throughput proteomic data[45].

While two-group comparison is one of the most useful designs, many studies involve a more complicated design structure, involving multiple groups, a block structure and/or time course measurements. MEACA provides a framework that is potentially generalizable to these designs with an extended mixed effects model and a modified set-level test statistic. It is our current work to adapt our approach to be applicable to analytical needs beyond two-group comparison.

## Methods

### The MEACA Method

We consider a gene expression (e.g. RNA-Seq or microarray) experiment, in which we compare the expression data of samples from two groups: a treatment group with *n*_1_ samples referred to as “cases” and a control group with *n*_2_ samples referred to as “controls” (*n*_1_, *n*_2_ ≥ 3). Suppose the expression levels of a set of m genes are observed for each sample. An unknown subset of these genes are differentially expressed between cases and controls, with varying sign and magnitude of DE effects. The genes are also allowed to have (negatively or positively) correlated expression levels. Our goal is to test whether a pre-defined set of test genes are enriched with DE signals compared to the background genes (i.e. genes not in the test set). The rest of this paragraph will provide a brief overview of the model underlying MEACA, with some technical details to be explained later. We will use a gene-level test statistic, denoted by *U_i_*, to capture the unknown DE signal of gene *i*. Let ***G*** be an m-dimensional vector defining the gene set of interest, where *G_i_* = 1 if and only if gene i is in the test set and *G_i_* = 0 otherwise (for any given gene set ***G*** is known). In the following sections, we will derive a model for *U_i_*’s conditional on ***G***, using a mixed-effects framework of the form (details to be explained later)

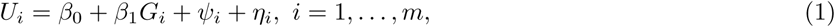

where *β*_1_ is a fixed effect capturing the mean difference between the test set and the background set, and *ψ*_*i*_ and *η*_*i*_ are random effects. The term *ψ*_*i*_ captures the variability among *U_i_*’s due to some genes being differentially expressed and some not, and to the varying magnitude of the DE effects. The variance of *ψ*_*i*_ depends on whether *G_i_* = 0 or 1, which allows the spread of gene-level statistics to be different between the test set and the background set. The *η*_*i*_’s account for the variability in *U_i_*’s due to sample-level noise and are allowed to be correlated with each other to accommodate inter-gene covariation.

To justify model (1) and to specify the modeling assumptions on *ψ*_*i*_ and *η*_*i*_, we will start by constructing a hierarchical model for the observed gene expression data, from which we will then derive a mixed-effects model for the gene-level statistics jointly for all the genes. Based on this model, we will then present our enrichment testing method, and discuss its connections with CAMERA.

#### A hierarchical model for gene expression data

We will start by presenting the hierarchical model for the observed gene expression data jointly for all genes, which will incorporate the following features. Firstly, for a given sample, the expression levels of different genes are allowed to be correlated. We further assume that the correlation structure is the same across samples. Secondly, different genes may have different baseline expression levels, where “baseline” refers to the average among controls. Thirdly, for any given gene, its mean expression level in the treatment group can be either higher, lower or the same compared to the control group, depending on whether the gene is up-regulated, down-regulated, or not differentially expressed. For the genes that are differentially expressed, their DE effects are modeled additively and are allowed to have heterogeneous signs and magnitudes. Finally, given a gene and its DE effect, the expression level is allowed to vary independently across samples, which captures measurement error and sample-level variability.

To present our model formally, we first introduce some notation. Let *n* = *n*_1_ + *n*_2_be the total sample size. Let ***X*** be an *n*-dimensional known vector of 1’s and 0’s denoting the case-control membership of the samples, with *X_i_* = 1 for a case and *X_i_* = 0 for a control. Let ***Y*** be an *m* by *n* matrix representing the expression data, in which each column is the expression profile for a sample and *Y_ij_*(1 ≤ *i* ≤ *m*, 1 ≤ *j* ≤ *n*) is the expression level of sample *j* at gene *i*. Let *μ*_*i*_ (1 ≤ *i* ≤ *m*) be the baseline expression level for gene *i*. The quantities *μ*_*i*_’s are treated as nuisance parameters and as we will see later do not contribute to our analysis. Let **Δ** = (Δ_1_, ⋯, Δ_*m*_) ^*T*^ be a vector for the additive DE effects for the genes. Gene *i* is not differentially expressed if Δ_*i*_ = 0, up-regulated if Δ_*i*_ > 0 and down-regulated if Δ_*i*_ < 0. We model **Δ** as a random effect, for which we will detail our assumptions later. Given *μ*_*i*_ and Δ_*i*_, the mean expression level for the control group and the treatment group are *μ*_*i*_ and *μ* _*i*_ + Δ_*i*_, respectively. Given these means, the noise in the observed expression data for the *j^th^* sample is denoted by the error vector ***ε***_*j*_ = (*ε*_*1j*_, ⋯, *ε*_*mj*_) ^*T*^, 1 ≤ *j* ≤ *n*. We assume ***ε***:= (***ε***_1_, ⋯, ***ε***_*m*_) to be independent of **Δ** and to have mean zero. Without loss of generality, we also assume Var(*ε*_*ij*_) = 1 for all genes and samples. For a real gene expression data set typically not satisfying this assumption, we can standardize the data by each gene to ensure that its empirical variance equals one before implementing our method (Additional file 1, Section A). For the covariance structure of ***ε***, we assume independence across samples and allow correlations between genes, namely

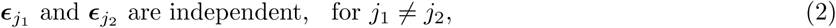

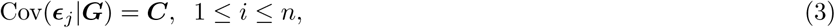

where ***C*** is an *m* by *m* inter-gene correlation matrix shared by all samples and is generally unknown. Putting these elements together, we obtain the following model for the expression data ***Y*** given ***X***

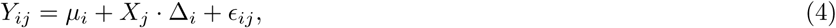

for 1 ≤ *i* ≤ *m*, 1 ≤ *j* ≤ *n*. The gene-set membership vector ***G*** enters this model via Δ_*i*_ and possibly *μ*_*i*_

#### Assumptions on the DE effects

Conditional on ***G***, we assume that the Δ_*i*_’s are mutually independent and come from either of the two distributions, 𝒟_1_ for the background set (i.e, *G_i_* = 0) and *𝒟*_2_ for the test set (i.e, *G_i_* = 1). We denote the expected values of *𝒟*_1_ and *𝒟*_2_ by *β*_0_ and *β*_0_ + *β*_1_, respectively, and their variances by *σ*_1_^2^ and *σ*_2_^2^, respectively. It follows that

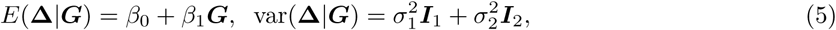

where ***I***_1_ and ***I***_2_ are diagonal matrices of dimension *m* with 0’s and 1’s on their diagonals. The 1’s in the diagonal of ***I***_1_ correspond to the genes with *G_i_* = 0 and those for ***I***_2_ to the genes with *G_i_* = 1.

Aside from the conditions in equation (5) on the first two moments, we do not impose on the DE effects, **Δ**, any specific distributional assumptions such as normality. For example, the distribution of a given Δ_*i*_ can put a positive probability mass on zero, which allows for the highly likely scenario in which some of the genes are not differentially expressed. To further illustrate our general framework for **Δ**, we present a simple model included by equation (5) as a special case. Suppose the *m* genes are independently sampled to be either differentially expressed or not. The probability for gene *i* to be differentially expressed is *p_t_* if *G_i_* = 1, or *p_b_* if *G_i_* = 0. For differentially expressed genes, their DE effects are sampled independently from a common distribution with mean *μ_δ_* and variance *σ_δ_^2^* Under these assumptions,

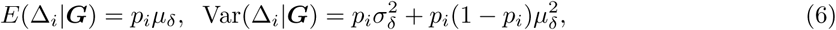

where *p_i_* = *p_t_* if *G_i_* = 1 and *p_i_* = *p_b_* if *G_i_* = 0 (Additional file 1, Section B). It can be shown that this model is a special case of equation (5), where *β*_1_ = 0 is equivalent to *p_b_* = *p_t_*.

#### Model for gene-level statistics

For each gene *i*, we consider the gene-level statistic *U_t_* given by

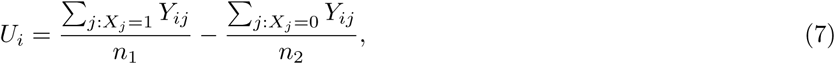

which is sample mean difference in the expression levels between cases and controls. Given our assumption that the expression data ***Y*** have been standardized so that *ε*_*i*_ has variance 1, *U_i_* is equivalent to the two-sample t-test statistic and provides a DE metric for gene *i*.

We will construct a quasi-likelihood model for the conditional distribution of ***U*** = (*U*_1_, ⋯ *U*_m_) *^T^* given ***G***, by deriving the conditional mean and covariance structures of ***U*** from the model for ***Y*** described in the previous two subsections. We first observe that combining equations (4) and (7) yields

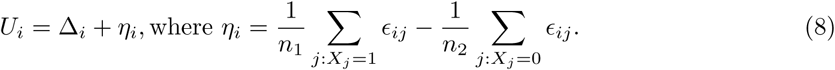

It can be shown (Additional file 1, Section C) based on equations (2), (3), (5) and (8) that

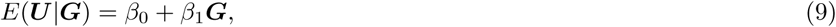

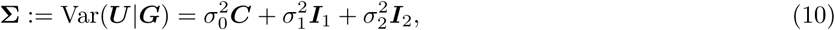

where *σ*_0_^2^ = 1/*n*_1_ + 1/*n*_2_ is a known parameter. We note that in equation (10), the covariance structure of ***U*** has three components, a component with ***C*** which accounts for the contribution of sample-level noise ***ϵ***, and two additional components from the heterogeneity of the DE effects **Δ**. It is noteworthy that both the ***C*** component and the **Δ** components contribute to the variance of *U_i_*’s, whereas only the ***C*** component contributes to the covariance between two *U_i_*’s. As the result, the correlation between two *U_i_*’s is affected by both the **Δ** components as well as the ***C*** component, with the former serving to increase the variance and therefore dilute the correlation. Ignoring the contribution of the former, as is done by some previously proposed methods including CAMERA, tends to lead to overestimation of the extent of inter-gene correlations for the *U_t_*’s.

Finally, we note that by letting Δ_*i*_ = *β*_0_ + *β*_1_*G_i_* + *ψ*_*i*_, equation (8) is equivalent to model (1) whose mean and variance are given by equations (9) and (10). The random effects *ψ*_*i*_’s capture the heterogeneity of the DE effects that are conditional on whether gene *i* belongs to the test set (*G_i_* = 1) or not (*G_i_* = 0).

#### The MEACA set-level test statistic

To detect patterns of the DE signals in the gene set of interest that stand out compared with genes not in the set, we test *H*_0_: 𝒟_1_ = 𝒟_2_ against *H*_1_: 𝒟_1_ ≠ 𝒟_2_. For example, for the special scenario given by equation (6), this amounts to testing *p_b_* = *p_t_* against *p_b_* ≠ *p_t_* To construct the set-level test statistic, we focus on the part of the alternative space where *E*(𝒟_1_) ≠ *E*(𝒟_2_), or equivalently *β*_1_ ≠ 0. We first consider the less interesting case with uncorrelated genes, in which ***C*** equals ***I***, an *m*-dimensional identity matrix. Under the quasi-likelihood model for ***U*** given in equations (9) and (10), the quasi-score statistic for *β*_1_ has the form *S* ∝ ***G***^*T*^ (***U*** − *β*̂_0_**1**_*m*_), where *β*̂_0_ = ***U***̅ is an estimate for *β*_0_ and **1**_*m*_ is a *m*-dimensional vector of 1’s. To perform a quasi-score test, one would divide *S*^2^ by its estimated variance under *H*_0_ and the assumption that ***C*** = ***I***. The resulting test statistic is

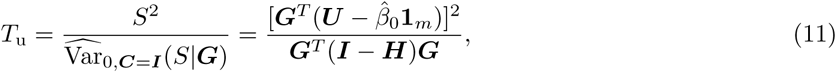

where 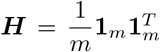 and the subscript “u” stands for “uncorrelated genes.” For the case of interest when inter-gene correlation is present, ***C*** is a non-trivial correlation matrix. We will again form our test statistic based on *S*. However, for the denominator of the statistic, the null variance of *S* will be evaluated under the quasi-likelihood model with a non-trivial ***C***. By equation (10), the variance of *S* is given by Var(*S*|***G***) = ***G***^*T*^ (***I*** − ***H***) **∑** (***I*** − ***H***) ***G***. Note that *H*_0_: 𝒟_1_ = 𝒟_2_ implies *σ_1_^2^* = *σ_2_^2^* Thus, under *H*_0_, ∑ = Var_0_(***U*** | ***G***) = *σ_0_^2^**C*** + *σ_1_^2^**I***, where *σ*_0_^2^ = 1/*n*_1_ + 1/*n*_2_ is known and *σ_1_^2^* is an unknown parameter. To estimate *σ_1_^2^* under *H*_0_, we observe that Var_0_(*U_i_*) = *σ_0_^2^* + *σ_1_^2^* and thus use 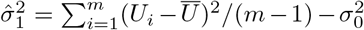. Therefore, assuming ***C*** is known, we can obtain the two-sided MEACA test statistic given by

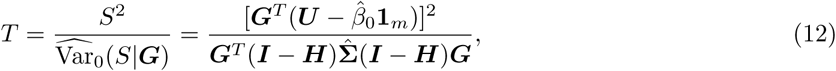

where 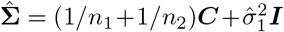 is a null estimate of **∑** and *β*̂_0_ = ***U***̄. Under suitable regularity conditions, significance of the test could then be assessed by comparing *T* to a *χ_1_^2^* distribution.

When it is desirable to test the one-sided alternative hypothesis that *E* (𝒟_1_) *E* < (𝒟_2_), one may use the signed squared root of *T* given by

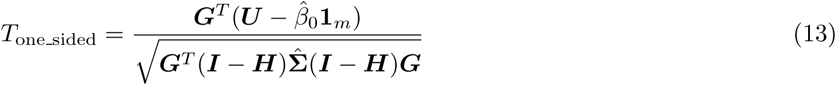

as the test statistic, whose *p*-value can be obtained by comparing to the standard normal distribution.

#### Estimating the inter-gene correlation matrix *C*

In practice, the inter-gene correlations are usually unknown. Therefore we substitute ***C*** with ***C***̂, the empirical correlation matrix of the expression data after possible DE effects are controlled for by centering the expression levels of cases and controls separately around zero. Formally, ***C***̂ is given 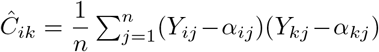, where 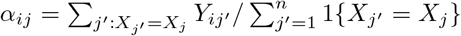 is the average expression level at gene *i* for all samples from the same group (either cases or controls) as sample *j*. In real data sets, the number of genes, *m*, is usually much greater than the sample size *n*, in which case ***C*** is a high-dimensional parameter that cannot be efficiently estimated by ***C***̂. Interestingly, however, we find that the MEACA test statistic *T* relies not on the entry-wise accurate estimation of ***C***, but only on three parameters involving the entries of ***C***, which can be much more realistically estimated given a moderate sample size. To demonstrate this, let *m*_1_ and *m*_2_ be the sizes of the test set and the background set, respectively (*m*_1_ + *m*_2_ = *m*). Also let *ρ*_1_ be the average correlation between two genes in the test set, *ρ*_2_ be the average correlation between two background genes, and *ρ*_3_ be the average correlation between a test gene and a background gene. Then, *ρ*_1_ is the mean of the off-diagonal entries in the *m*_1_ × *m*_1_ sub-matrix of ***C*** made up of rows and columns corresponding to the test set, *ρ*_2_ is that in the *m*_2_ × *m*_2_ sub-matrix corresponding to the background set, and *ρ*_3_ is the mean of the entries in the *m*_1_ × *m*_2_ sub-matrix of ***C*** corresponding to the cross-covariance between the test and the background sets. It can be shown that the denominator of the MEACA test statistic given in equation (12) can be written as

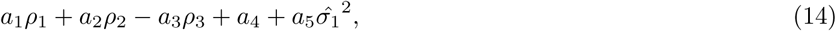

where *a*_1_, ⋯, *a*_5_ > 0 are constants that do not depend on ***C*** (for details see Additional file 1, Section D). Therefore, the MEACA test statistic depends on ***C*** only through *ρ*_1_, *ρ*_2_ and *ρ*_3_.

#### Connections with CAMERA

Model (1) and expression (14) also help reveal the connections between CAMERA and our method. When considered under our framework with Var(ϵ_*ij*_) = 1 and equation (7) as the gene-level statistics, the CAMERA approach can be viewed as a score test derived from a model which effectively assumes (A1) and(A2) introduced in Results, which are further explained as follows:

(A1) The random effect *ψ*_*i*_ =0 can be dropped from model (1) for both genes in the test set and those in the background set. Or equivalently, *σ_1_^2^* = *σ_2_^2^* = 0 in equation (10). This amounts to assuming, both in the test set and in the background set, that either none of the genes are differentially expressed or all genes are differentially expressed with the exact same DE effect;
(A2) The inter-gene correlation structure satisfies *ρ*_2_ = *ρ*_3_ = 0, which means that inter-gene correlations are present only among genes in the test set, not among background genes or between background and test genes.

Both assumptions are likely violated in reality. In particular, it is likely for both the test set and the background set that some genes are differentially expressed while others are not, and that the genes that are differentially expressed vary in terms of the signs and magnitudes of their DE effects. In our model, this is accounted for by a non-trivial *ψ*_*i*_ term or equivalently by the heterogeneity in the Δ_*i*_’s, which adds to the variances of ***U***_*i*_’s without contributing to their pairwise covariances. However, with Assumption (A1), CAMERA effectively ignores the Δ_*i*_ heterogeneity and consequently under-estimates the variances of ***U***_*i*_’s and over-estimates the correlations between ***U***_*i*_’s. This tends to result in over-adjustment of inter-gene correlations in enrichment testing and lead to conservative type 1 error and power loss. In the setup given by equation (14), this issue would be reflected by incorrectly calculated constants *a*_1_⋯, *a_5_* which overall would produce a greater than necessary denominator in the set-level test statistic and thus tends to drive the *p*-value towards the non-significant side. With Assumption (A2), ignoring a positive *ρ*_2_ has the effect of under-estimating the null variance of the set-level test statistic and thus may inflate type 1 error, whereas ignoring a positive *ρ*_3_ has the opposite effect. Overall, whether CAMERA results in a conservative or anti-conservative type 1 error will depend on how these factors act upon each other. In simulation studies, we will explore how CAMERA behaves in different scenarios.

### Simulation Study Design

In this section, we will specify the setup of our type 1 error and power simulation studies. Let ***Y***_*j*_ be a vector denoting the expression profile of sample *j*. Conditional on the genes’ DE effects, we simulate the ***Y***_*j*_’s independently from a multivariate normal distribution with unit variance and **ρ*_i_1_, i_2__* = Cor(*Y_i_1_, j_, Y_i_2_, j_*) as the correlation coefficient between genes *i*_1_ and *i*_2_. We assume a common pairwise correlation coefficient for genes from the same category (either the test set or the background set): Cor(*Y_i_1__, Y_i_2__*) = **ρ*_1_* if genes *i*_1_ and *i*_2_ are both test genes (i.e., *G_i_1__* = 1, *G_i_2__* = 0), Cor(*Y_i_1__, Y_i_2__*) = **ρ*_2_* if they are both background genes (i.e., *G_i_1__* = 1, *G_i_2__* = 0). For a test gene and a background gene (i.e., *G_i_1__* = 1, *G_i_2__* = 0), we assume Cor(*Y_i_1__, Y_i_2__*) = *ρ*_3_. We examine five different correlation structures, as listed in Results.

The simulations run as follows. First, we consider a total of *m* = 500 genes, of which *m*_1_ = 100 genes are in the test set and the remaining *m*_2_ = 400 genes in the background set. Second, we randomly sample genes to be differentially expressed with probability *p_t_* in the test set and with probability *p_b_* in the background set. If gene *i* is sampled to be differentially expressed, we simulate its DE effect Δ_*i*_ from a normal distribution *N*(2,1), and if gene *i* is not differentially expressed, we set Δ_*i*_ = 0. Third, we set the mean expression levels of the *m* genes to be ***μ***_1_ = **0**_*m*_for a control sample and ***μ***_2_ = **Δ** for a case sample. Fourth, for each of the *n*_1_ = 25 samples in the control group, we simulate its expression profile independently from a multivariate normal distribution MVN(***μ***_1_, **∑**), where **∑** = [Cov(*Y_i_1__, Y_i_2__*)]_*m × m*_ is the covariance matrix corresponding to one of the structures in (a)-(e) detailed in the previous paragraph. For each of the *n*_2_ = 25 samples in the treatment group, we simulate its expression profile from MVN(***μ***_2_, **∑**).

Further assumptions on *p_t_* and *p_b_* will complete our generating model used in the type 1 error and power simulations. Table 1 summarizes the configurations of *p_b_* and *p_t_* we consider. In order to examine how the presence of DE and the heterogeneity of the DE effects may affect various enrichment tests, for each correlation structure in (a)-(e), we conduct two groups of simulations: genes in the background set are allowed to be differentially expressed in group II but not in group I (so Assumption (A1) holds for group I but not for II). In both type 1 error and power simulations, we set the DE probability for the background genes to be *p_b_* = 0% in group I and *p_b_* = 10% in group II. In the type 1 error simulations, we have *p_t_* = *p_b_* under the null. In the power simulations, we consider four different scenarios, *S*_1_— *S*_4_, for the alternative hypothesis corresponding to different levels of enrichment: for genes in the test set, we set the DE probability to be *p_t_* = 5% (*S*_1_), 10% (*S*_2_), 15% (*S*_3_) and 20% (*S*_4_) in group I, and 15% (*S*_1_), 20% (*S*_2_), 25% (*S*_3_) and 30% (*S*_4_) in group II.

### Software Implementation of Methods

To implement MEACA, we have developed an publicly available R package. sig-Pathway [10], MRGSE [26], CAMERA [11] are all implemented in the limma [46] package of the Bioconductor [47] project. QuSAGE [12] is available as an R package under the same name. GSEA [14] is implemented in the R-GSEA script (http://software.broadinstitute.org/gsea/index.jsp). We note however, that GSEA can yield p values that are exactly zero, which have been shown to be inaccurate for permutation tests [48]. To avoid exactly zero *p*-values, we follow the recommendation of Phipson et al. [48] and calculate the GSEA *p* value using (*b* + 1)/(*K* + 1), where *K* is the total number of permutations performed and *b* out of the *K* permutations result in statistics that are more extreme than the observed statistic. In the simulation studies, we use default of the R-GSEA program *K* = 999. In the real data analysis, where we increase the number of permutations, *K*, to 9999 due to the need to more accurately estimate smaller *p*-values.

## Declarations

### Acknowledgements

We thank Yanming Di, Sarah Emerson and Wanli Zhang for helpful discussion in preparing this manuscript. We thank Dr. Adam Labadorf for providing information about the HD gene expression data.

### Availability of data and material

MEACA is implemented in an R package available at http://www.science.oregonstate.edu/~jiangd/. The HD data were downloaded from NCBI GEA GSE64810 (http://www.ncbi.nlm.nih.gov/geo/). The lymphoblastoid cells data were downloaded from broad institute (http://software.broadinstitute.org/gsea/datasets.jsp).

### Authors’ contributions

DJ conceived of and designed the research. BZ performed the experiments and analyzed the data. DJ and BZ wrote the manuscript. Both authors read and approved the final manuscript.

### Ethics approval and consent to participate

Not applicable.

### Consent for publication

Not applicable.

## Additional Files

Additional file 1 (pdf) **Supplemental Method**. A. Standardization; B. Special case of model (6); C. Covariance matrix for *U_i_*’s, the gene-level test statistics; D. Constants in equation (14).

